# COX6B1 secures a redox-sensitive step in early cytochrome *c* oxidase assembly

**DOI:** 10.1101/2025.03.25.645161

**Authors:** Kristýna Čunátová, Marek Vrbacký, Michal Knězů, Alena Pecinová, Lukáš Alán, Josef Houštěk, Erika Fernández-Vizarra, Tomáš Mráček, Petr Pecina

## Abstract

COX6B1 is a nuclear-encoded subunit of the human mitochondrial cytochrome c oxidase (cIV) located in its intermembrane space-facing region. The relevance of COX6B1 in mitochondrial physiopathology was highlighted by the missense pathogenic variants associated with cIV deficiency. Despite the assigned COX6B1 role as a late incorporation subunit, the COX6B1 human cell line knock-out (KO) exhibited a total loss of cIV. To get a deeper insight into the mechanisms driving the lack of cIV assembly or destabilization in the absence of COX6B1, we used the COX6B1 KO cell background to express alternative oxidase and COX6B1 pathogenic variants. These analyses uncovered that the COX6B1 subunit is indispensable for redox-sensitive early cIV assembly steps, besides its contribution to the stabilization of cIV in the late assembly stages. In addition, we have evidenced the incorporation of partially assembled cIV modules directly into supercomplex structures, supporting the ‘cooperative assembly’ model for respiratory chain biogenesis.

## 1 Introduction

Human cytochrome *c* oxidase (cIV, COX) is the terminal component and pacemaker of the mitochondrial respiratory chain (RC) ^1^. cIV is found in the inner mitochondrial membrane in diverse forms – monomeric, dimeric (IV_2_), and also participates in supramolecular structures interacting with other RC complexes I (cI) and dimeric III (cIII_2_), forming the so-called supercomplexes (SCs, typically SC III_2_IV, and respirasome - SC I III_2_IV) ^2^. Human monomeric cIV is a protomer of three mitochondrial DNA (mtDNA)-encoded subunits (MT-CO1, MT-CO2, and MT-CO3) constituting its catalytic core, and eleven supernumerary nuclear (nDNA)-encoded subunits (COX4, COX5A, COX5B, COX6A, COX6B, COX6C, COX7A, COX7B, COX7C, COX8, and NDUFA4, alias COXFA4) ^3–5^, which are necessary for cIV assembly and maintenance. Seven of the nDNA-encoded subunits (COX4, COX6A, COX6B, COX7A, COX7B, COX8, and NDUFA4) exist in more than one variant, so-called isoforms, which are encoded by distinct genes ^6,7^ and facilitate adaptation of cIV function in specific conditions and tissues ^8,9^. Even if there are these tissue-specific differences in composition, cIV structure and subunit number is consistent ^8,10–13^. In addition, the association of cIV within RC SCs might help fine-tune its function to specific conditions ^8^.

The current ‘modular model’ of *de novo* cIV assembly implies individual modules (MT-CO1, MT-CO2 and MT-CO3) – each one formed by an mtDNA-encoded subunit accompanied by some of the nDNA-encoded subunits – that assemble in a process secured by a group of specific chaperons and assembly factors (AFs), all of them contributing towards the formation of a fully active cIV (Figure 1A) ^14–16^. The incorporation of late-stage subunits, belonging to the MT-CO3 module, such as COX6A, COX6B, and COX7A, has been observed to happen both during *de novo* cIV biogenesis, but also as replacement of the components of previously assembled cIV structures ^17^. Moreover, several studies have presented that even incomplete cIV may attach to and stabilize SC I III_2_ in the absence of a fully assembled cIV ^12,18–21^. This indicates that parallel pathways of cIV assembly occur to form the monomeric cIV and the respirasome ^19^. These observations are consistent with the proposed ‘cooperative assembly’ model, in which SC biogenesis is not the product of the association of fully assembled RC complexes, but instead is the product of joining partially assembled modules of each component complex ^22^.

**Figure 1:**
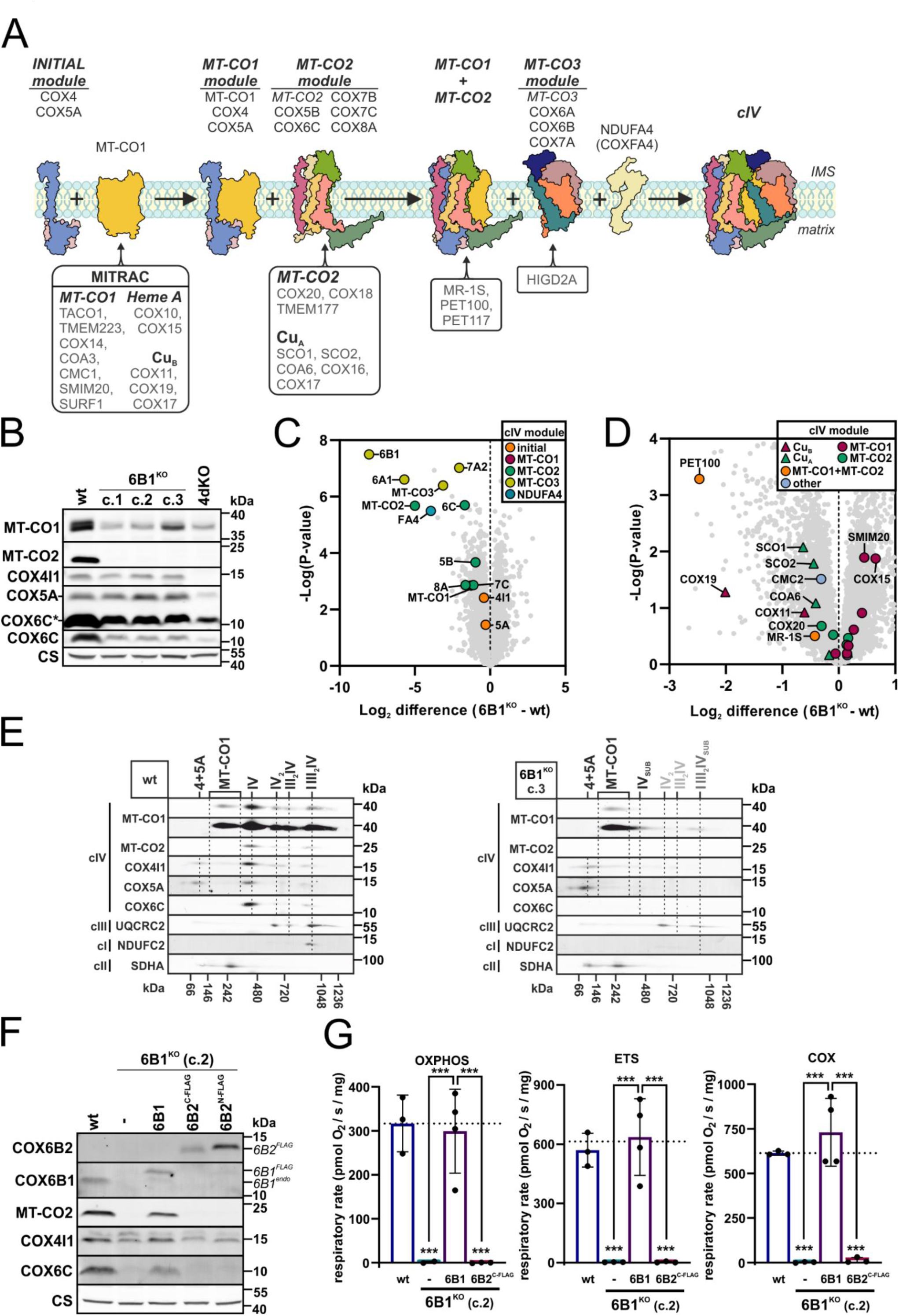
COX6B subunit is indispensable for early human cIV assembly and function. (A) Modular model of monomeric cIV assembly. Initially, the MT-CO1 module is formed by the connection of the initiating COX4-COX5A module with MT-CO1, which maturates in the MITRAC complex (mitochondrial translation regulation assembly intermediates of cytochrome *c* oxidase) ^14,63,64^. The second module consists of MT-CO2, which undergoes copper insertion, to form the binuclear Cu*_A_* site, assisted by SCO1, SCO2, COX16, COX17, and COA6 chaperones, and subunits COX5B, COX6C, COX7B, and COX8A ^38,65–68^. In the end, the third module containing MT-CO3, COX6A, COX6B, and COX7A subunits joins the nascent cIV under the assistance of HIGD2A, and the addition of NDUFA4 finalizes the assembly of complete and fully functional cIV ^12,15,33^. cIV subunits are depicted in various colors and mentioned on the top, assembly factors (AFs) of individual modules are noted on the bottom. Cartoons are based on human cIV cryo-EM structure (PDB ID 5Z62) ^62^. (B) Representative SDS-PAGE/WB analysis of the protein steady-state level of cIV subunits (MT-CO1, MT-CO2, COX4I1, COX5A, COX6C) and citrate synthase (CS) as the loading control in whole cell-lysates. COX6C* represents an overexposed signal of COX6C from the same image. See also Figure S1A. (C) Differential content of cIV subunits between wt and 6B1^KO^ cells. Volcano plot represents LFQ-MS analysis (wt: n = 4; 6B1^KO^ combines c.1, c.2 and c.3: n = 2 per each / n = 6 in total) of all analyzed proteins (grey), and subunits of cIV modules (initial: orange, MT-CO1: magenta, MT-CO2: green, MT-CO3: yellow, NDUFA4: blue). COX6B1 protein missing in 6B1^KO^ was visualized thanks to the imputation step (missing values replaced from normal distribution) performed during the Perseus analysis of the LFQ-MS data. For individual 6B1^KO^ clones see Figure S1B. (D) Differential content of cIV assembly factors (AFs) between wt and 6B1^KO^ cells. Volcano plot represents LFQ-MS analysis (wt: n = 4; 6B1^KO^ combines c.1, c.2 and c.3: n = 2 per each / n = 6 in total) of all analyzed proteins (grey), and cIV assembly factors (MT-CO1 metalation: triangle in magenta, MT-CO1 maturation: circle in magenta, MT-CO2 metalation: triangle in green, MT-CO2 maturation: circle in green, MT-CO1+MT-CO2 metalation: triangle in yellow, MT-CO1+MT-CO2 association: circle in orange, other: circle in blue). For individual 6B1^KO^ clones see Figure S1C. (E) 2D (BN/SDS)-PAGE/WB detection of cIV (MT-CO1, MT-CO2, COX4I1, COX5A, and COX6C antibodies), cIII (UQCRC2 antibody), and cI (NDUFC2 antibody) in wt (left) and 6B1^KO^ (c.3, right) mitochondrial fraction. Antibody against cII (SDHA) was used as a loading control. (F) Representative SDS-PAGE/WB analysis of the protein steady-state level of cIV subunits (COX6B2, COX6B1, MT-CO2, COX4I1 and COX6C) and CS as the loading control in whole cell-lysates. FLAG-tagged COX6B1 and COX6B2 proteins are marked by 6B1^FLAG^ and 6B2^FLAG^ respectively, endogenous COX6B1 is labelled with 6B1^endo^ (in italics). See also Figure S1D. (G) Respiratory rates in OXPHOS, ETS, and COX states plotted as the mean ± S.D. value of wt (n = 3), 6B1^KO^ (n = 2), 6B1^KO^+6B1 (n = 4), and 6B1^KO^+6B2^C-FLAG^ (n = 3). One-way ANOVA (*** p < 0.001) was performed. See also Figure S1E.

COX6B is a nDNA-encoded subunit associated to cIV facing the intermembrane space; it was thought to stabilize the dimeric cIV_2_ structure and support the formation of the binding site for cytochrome *c* ^4^. While the ubiquitous mammalian isoform COX6B1 is present in all cell types and tissues, expression of the COX6B2 isoform is restricted to testes ^23^. The functional importance of this subunit is underscored by the existing link between decreased COX6B2 content and altered sperm motility and decreased fertility ^24,25^. In addition, COX6B2 expression in other tissues might be associated with carcinogenesis ^26–28^.

Even though it is a peripherally associated subunit, COX6B1 is important for cIV structural stability/assembly or function - *COX6B1* missense pathogenic variants have been identified as the genetic cause of childhood-onset mitochondrial disease ^29–31^. The first two identified variants R20C and R20H led to a destabilization of COX6B1 interaction with cIV, and decreased cIV activity. Similarly, the knock-down of COX6B1 expression manifested as cIV deficiency ^29^. Therefore, based on multiple observations, COX6B subunit was perceived as one of the subunits joining cIV assembly in the very late stages ^12,14,32,33^. In contrast to this idea, we found that the knock-out (KO) of COX6B1 in human cells, produced a total loss of cIV ^20^, indicating an essential but largely unexplored role for COX6B1 in the early biogenesis of cIV.

Here, we have utilized *COX6B1* KO cells and subsequent models prepared by expression of its isoforms, its pathogenic variants (R20C, R20H) or alternative oxidase (AOX), which has allowed us to clarify the role of the COX6B1 subunit in the life cycle of cIV.

## 2 Results

### 2.1 COX6B subunit is indispensable for early human cIV assembly and function

We decided to generate COX6B1 KO as a model of late-stage cIV assembly disruption ^14,16^ (Figure 1A), which would contrast with the harsh COX4 KO models we generated previously ^20,34^. For this purpose, we utilized CRISPR to KO the *COX6B1* gene in HEK293 cells and identified a positive clone (6B1^KO^, clone 2), which was characterized previously ^20^. In the present work, we included another two 6B1^KO^ clones (clone 1, clone 3) to exclude clonal specificity and elucidate the reason for the severe cIV deficiency phenotype in COX6B1-lacking cells ^20^. Analysis of the steady-state level of cIV subunits in the three 6B1^KO^ clones (c.1, c.2, c.3) by SDS-PAGE/western blot (WB) (Figures 1B, S1A) consistently revealed the downregulation of numerous cIV subunits. MT-CO1 and COX6C subunits were significantly decreased compared to wild type (wt) cells, and MT-CO2 was completely undetectable in 6B1^KO^, as in the severe cIV deficiency model 4dKO (KO of the early assembly COX4 subunit) ^34^. On the contrary, the levels of COX4 and COX5A subunits in 6B1^KO^ were similar to wt, suggesting preservation of the early assembly intermediate of cIV (Figures 1B, S1A). This finding was confirmed and extended by whole proteome analysis of three 6B1^KO^ clones and the wt cells, using label-free quantification mass spectrometry (LFQ-MS). The levels of MT-CO3 module subunits (MT-CO3, COX6A1, COX7A2) and NDUFA4 were the most decreased, but MT-CO2 module subunits MT-CO2 and COX6C, were also significantly reduced in COX6B1 lacking cells (Figures 1C, S1B). The LFQ-MS data for cIV assembly factors (AFs) in the three 6B1^KO^ clones revealed a significant decrease of PET100, which is thought to secure the stability of MT-CO2 joining the MT-CO1 module ^14^ (Figures 1D, S1C). Apart from PET100, the most decreased AFs were those involved in MT-CO1 (COX19, COX11) and MT-CO2 (SCO1, SCO2, COA6) metalation (Figure 1D). Altogether, these results suggest a block in cIV assembly at the stage of MT-CO2 maturation or/and incorporation.

To inspect the *COX6B1* KO effect on the native cIV forms and its assembly intermediates, we performed a first dimension Blue-Native (BN-PAGE) followed by a denaturing (SDS-PAGE) second dimension and WB-immunodetection analysis. These experiments showed the total loss of fully assembled monomeric cIV (IV), cIV dimer (IV_2_) as well as SC III_2_IV in 6B1^KO^. Nevertheless, several cIV assembly intermediates were accumulated in 6B1^KO^ clones in comparison to wt (Figures 1E - clone 3, cf. also 4C - clone 2, S3B - clone 1). The most prominent assembly intermediates in 6B1^KO^ cells, composed of subunits MT-CO1, COX4 and COX5A, correspond to MT-CO1 module, the initial intermediate (COX4+COX5A), and their association, preceding the incorporation of the MT-CO2 module. In addition, a signal corresponding to MT-CO1 could be detected in the higher molecular weight part of the blot (at ∼1MDa) in the 6B1^KO^ samples and co-migrating with cIII_2_ (Figure 1E). This implies a non-negligible presence of SC species formed by cI, the dimer of cIII and at least MT-CO1.

To test whether the *COX6B1* KO phenotype could be rescued not only by overexpression of COX6B1, but also by the COX6B2 isoform, we stably expressed either COX6B1 or COX6B2 in the 6B1^KO^ clone 2 (Figure 1F). The complementation of 6B1^KO^ by a C-terminal FLAG-tagged COX6B1 (6B1^KO^+6B1) resulted in a significant recovery of the MT-CO2 and COX6C levels (Figure 1F, S1D). This was paralleled by the restoration of respiratory capacities of COX as well as of the OXPHOS and ETS states to levels comparable to the wt (Figure 1G), details of the respirometry protocol are described in Figure S1E) On the other hand, the expression of either a C-terminal or N-terminal FLAG-tagged COX6B2 isoform failed to even marginally rescue either MT-CO2 levels (Figures 1F, S1D) or mitochondrial respiration (Figure 1G). This may indicate, that other cIV tissue-specific isoforms and/or other factors are required for COX6B2 incorporation into cIV. At the same time, it explains why COX6B2 upregulation cannot functionally complement pathologic COX6B1 variants in patients since the same is likely valid also for tissues other than testes.

The presented data indicate that the COX6B1 subunit is essential for proper cIV assembly and function, since its loss leads to total cIV deficiency caused by a block in cIV biogenesis at a stage subsequent to MT-CO1+COX4+COX5A subassembly formation.

### 2.2 R20C pathogenic variant of COX6B1 affects an early cIV assembly, unlike R20H that disrupts cIV stability

Interestingly, the cIV assembly defect observed in patients carrying homozygous pathogenic variants in COX6B1 is much milder than the defect in the 6B1^KO^ cells ^29^. Therefore, we next explored the effects of the R20H and R20C COX6B1 pathogenic variants (Figure 2A, 2B) when expressed in the HEK293 6B1^KO^ cell line. The expression levels of a C-terminal FLAG-tagged COX6B1 R20H variant (6B1^KO^+R20H) were comparable to those of the wt COX6B1 (6B1^KO^+6B1, Figure 2C, S2). In contrast, the overexpression of a C-terminal FLAG-tagged R20C variant (clone 2, to produce 6B1^KO^+R20C) resulted in much lower COX6B1 levels, most probably because of a decreased stability of the R20C variant (Figures 2C, S2). The levels of the MT-CO2 subunit were decreased in both 6B1^KO^+R20H and 6B1^KO^+R20C cells compared to 6B1^KO^+6B1, while COX6C content was the same in the three cell lines, being decreased relative to wt (Figures 2C, S2). COX4 subunit levels were the same in the five cell lines (wt, 6B1^KO^ and the three overexpression lines) (Figures 2C, S2).

**Figure 2:**
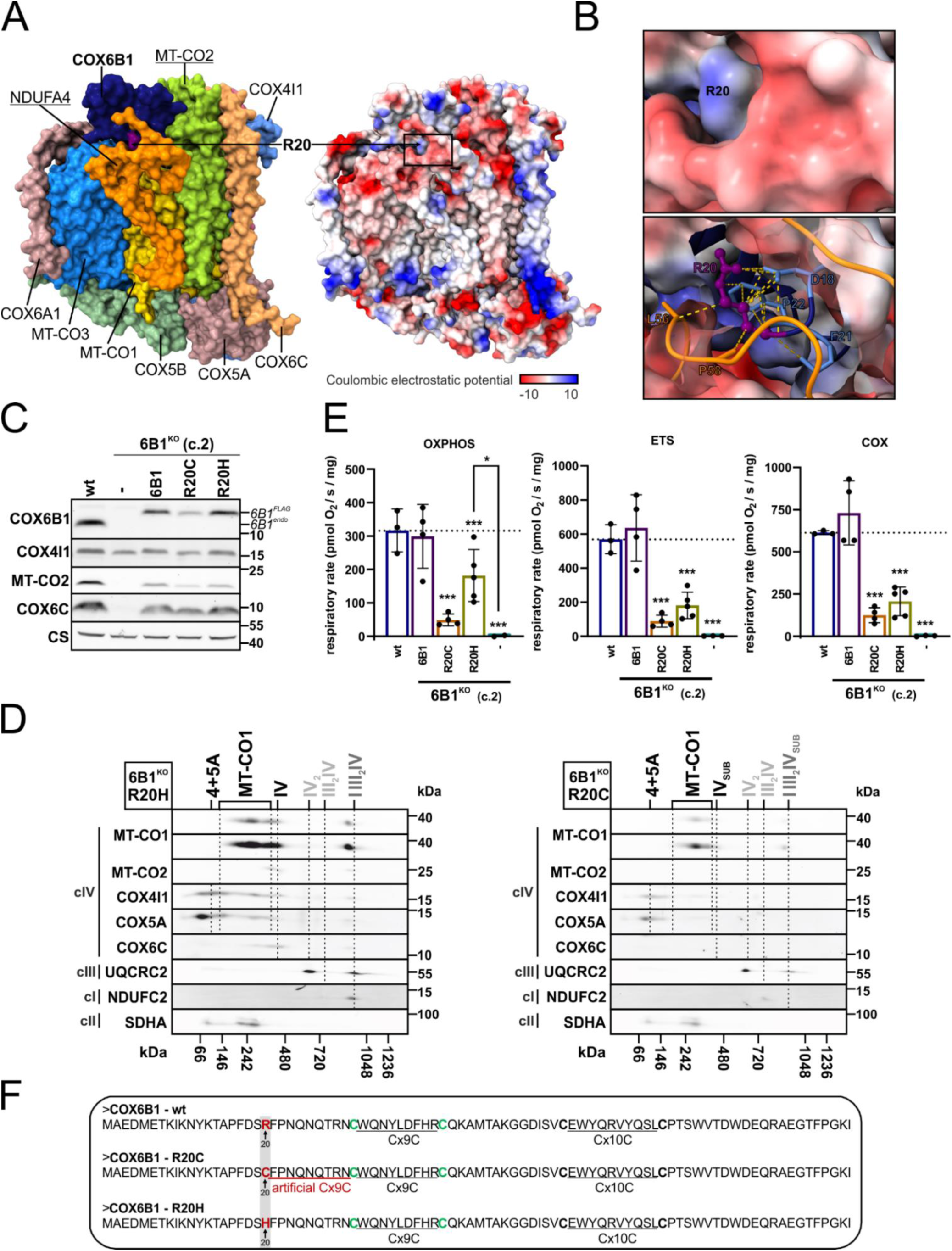
Characterization of R20C and R20H pathogenic variants of COX6B1 in HEK293 cellular models. (A) Position of COX6B1 R20 in human cIV structure. On the left: structural subunits of cIV are indicated in different colors, with emphasis on the location of R20 (violet) of COX6B1 (dark blue) in cryo-EM structure of human cIV (PDB 5Z62) ^62^. On the right: coulombic electrostatic potential visualization of cIV. The images were created using UCSF ChimeraX ^69,70^. (B) Detailed image of COX6B1 - R20 using coulombic electrostatic potential visualization, based on human cIV structure (PDB 5Z62). The images were created with UCSF ChimeraX ^69,70^. (C) Representative SDS-PAGE/WB analysis of the protein steady-state level of cIV subunits (COX6B1, COX4I1, MT-CO2, COX6C) and CS as the loading control, in whole cell-lysates. FLAG-tagged COX6B1 and endogenous COX6B1 proteins are marked with 6B1^FLAG^ and 6B1^endo^ (in italics) respectively. See also Figure S2. (D) 2D (BN/SDS)-PAGE/WB detection of cIV (MT-CO1, MT-CO2, COX4I1, COX5A, and COX6C antibodies), cIII (UQCRC2 antibody), and cI (NDUFC2 antibody) in 6B1^KO^+R20H (left) and 6B1^KO^+R20C (right) mitochondrial fraction. Antibody against cII (SDHA) was used as a loading control. (E) Respiratory rates in OXPHOS, ETS, and COX states plotted as the mean ± S.D. value of wt (n = 3), 6B1^KO^+6B1 (n = 4), 6B1^KO^+R20C (n = 4), 6B1^KO^+R20H (n = 5), and 6B1^KO^ (n = 2). One-way ANOVA (* p < 0.05, *** p < 0.001) was performed. Plotted data of wt, 6B1^KO^ and 6B1^KO^+6B1 originate from Figure 1G. (F) Primary sequence of wt, R20C and R20H variants of COX6B1 protein. The artificial Cx9C motif created by the R20C pathogenic variant is highlighted in red.

The cIV defect observed in cells expressing the two different R20 variants was not equivalent, as the R20C induced a harsher assembly defect with only little detectable amounts of the cIV monomer and undetectable cIV dimer (IV_2_) and SC III_2_IV (Figure 2D). Interestingly enough, MT-CO1 was detected in the respirasome in both R20C and R20H cell lines (Figure 2D). The milder cIV assembly defect was also reflected with higher levels of respiration (coupled – OXPHOS, uncoupled - ETS state and cIV capacity - COX) in the 6B1^KO^+R20H than in the 6B1^KO^+R20C, which despite the very low abundance of fully assembled cIV native forms could sustain detectable levels of oxygen consumption. Specifically, the respiration rates were approximately 2,5- and 3-times decreased in the ETS (maximal respiratory capacity after uncoupling) state in 6B1^KO^+R20H and 6B1^KO^+R20C, respectively, compared to wt and 6B1^KO^+6B1 cell lines (Figure 2E). Thus, introduction of R20C variant disrupted cIV native forms and respiration to a higher extent than R20H, similar to the observations in patient-derived fibroblasts ^29,30^. Interestingly, the additional cysteine residue in COX6B1-R20C variant forms an artificial Cx9C motif adjacent to the one in the COX6B1 wt primary sequence (Figure 2F). This is likely to interfere with proper folding within the IMS ^35^, resulting in decreased stability and lower steady-state levels of the protein (Figure 2C, S2).

To reveal the ability of R20C and R20H variants to associate with the native cIV forms, as well as their effect on cIV subunit and AF amounts, we used the mild detergent digitonin to solubilize the mitochondrial membranes and performed immunoprecipitation of cIV, followed by MS analysis of the co-immunopurified fractions (Figure 3). According to these analyses, the same cIV structural subunits were found reduced in 6B1^KO^+R20H, 6B1^KO^+R20C and 6B1^KO^ compared to wt but not to the same degree (Figure 3A). The initial module was least affected (COX4 and COX5A being unchanged) and MT-CO1 was within the less decreased subunits. MT-CO2 module subunits were, on average, underrepresented to a larger extent, and MT-CO3 plus NDUFA4 were the most affected (Figure 3A). In concordance with the previous results (Figures 2C, 2D), 6B1^KO^+R20C showed more profound decrease of subunits incorporated into the native cIV subassemblies, including COX6B1 (containing the variant), in comparison with 6B1^KO^+R20H (Figure 3A). Analysis of cIV bound AFs showed higher association of CMC1, COX14 and COA3, which stabilize MT-CO1 ahead of MT-CO2 incorporation and metalation, and lower abundance of PET100 in 6B1^KO^, but also in 6B1^KO^+R20H and 6B1^KO^+R20C compared to wt (Figure 3B). Strikingly, level of bound COA6, a factor responsible for copper delivery to MT-CO2 ^36–38^, was increased in the case of 6B1^KO^+R20H and 6B1^KO^+R20C models, while being decreased in 6B1^KO^ in comparison to wt (Figure 3B). This difference reinforces the possibility of the COX6B1 subunit having a relevance for MT-CO2 maturation. Further, the difference between R20C and R20H variants of COX6B1 was highlighted by diverse alteration of cIV bound AFs ensuring MT-CO1 and MT-CO2 metalation and maturation, which were diminished in 6B1^KO^+R20C relative to 6B1^KO^+R20H (Figure 3B).

**Figure 3:**
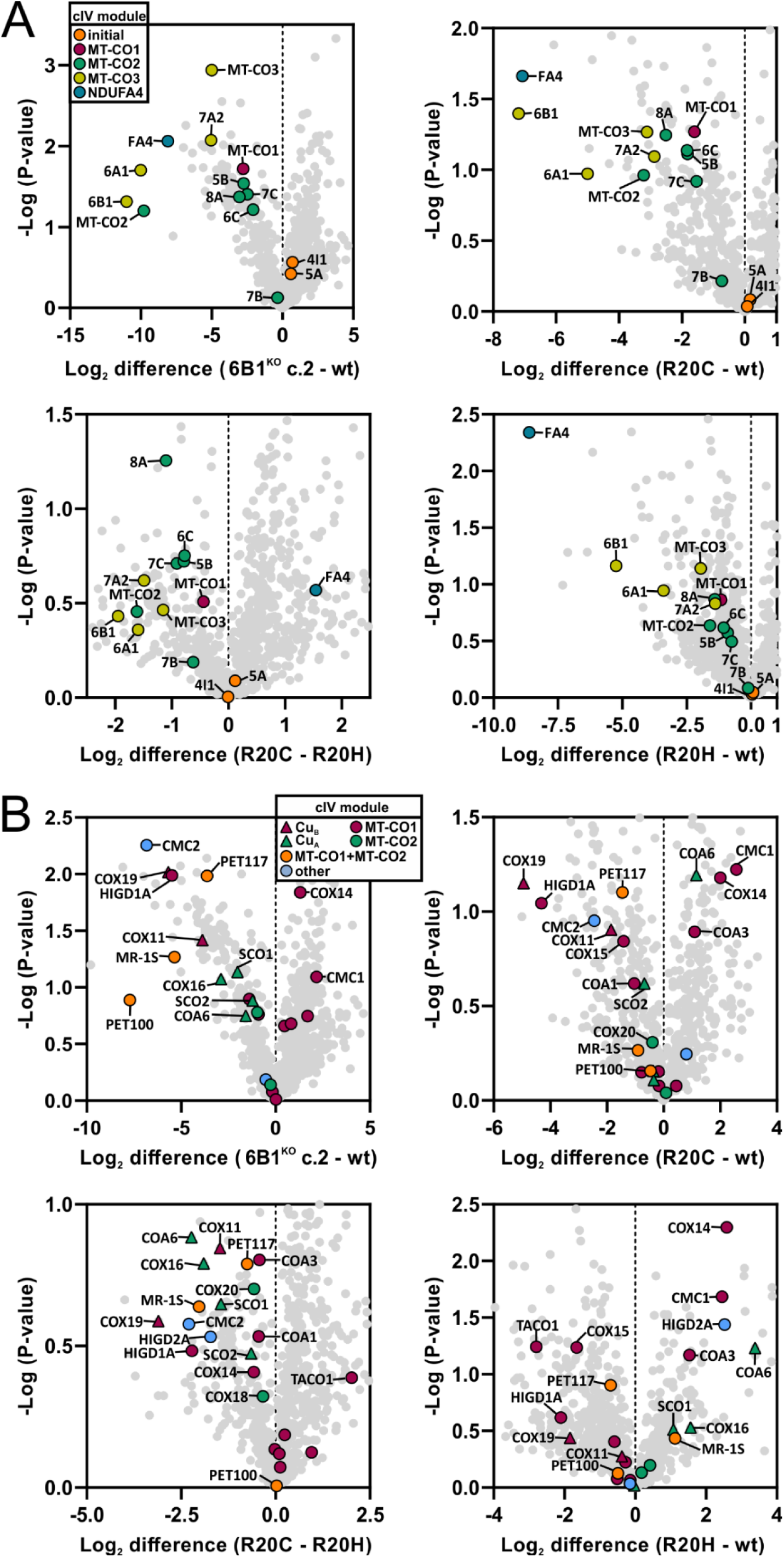
R20C pathogenic variant of COX6B1 affects an early cIV assembly, unlike R20H that disrupts cIV stability. (A) Differential amount of cIV subunits associated in cIV forms between wt and 6B1^KO^ (top left), 6B1^KO^+R20C (top right), 6B1^KO^+R20H (bottom right), and between 6B1^KO^+R20C and 6B1^KO^+R20H (bottom left). Volcano plots represent LFQ-MS analysis (n = 2) of all detected proteins (grey), and subunits of cIV modules (initial: orange, MT-CO1: magenta, MT-CO2: green, MT-CO3: yellow, NDUFA4: blue). COX6B1 protein missing in 6B1^KO^ was visualized thanks to the imputation step (missing values replaced from normal distribution) performed during the Perseus analysis of the LFQ-MS data. (B) Differential amount of cIV AFs associated with cIV forms between wt and 6B1^KO^ (top left), 6B1^KO^+R20C (top right), 6B1^KO^+R20H (bottom right), and between 6B1^KO^+R20C and 6B1^KO^+R20H (bottom left). Volcano plots represent LFQ-MS analysis (n = 2) of all detected proteins (grey), and cIV assembly factors (MT-CO1 metalation: triangle in magenta, MT-CO1 maturation: circle in magenta, MT-CO2 metalation: triangle in green, MT-CO2 maturation: circle in green, MT-CO1+MT-CO2 metalation: triangle in yellow, MT-CO1+MT-CO2 association: circle in orange, other: circle in blue).

### 2.3 Expression of alternative oxidase (AOX) restores cIV assembly with negligible cIV function in COX6B1 deficient cells

Our data indicate a block at the level of MT-CO2 maturation in 6B1^KO^ cells. Since MT-CO2 maturation is a redox-sensitive process ^39,40^ we modified the redox status in the 6B1^KO^ mitochondria via xenoexpression of an alternative oxidase (AOX), an enzyme with CoQ:O_2_ oxidoreductase activity present in the mitochondria of certain organisms ^20,41,42^. SDS-PAGE/WB analysis showed that stable expression of AOX in 6B1^KO^ (6B1^KO^+AOX, c.1 and c.2) led to a significant increase in the steady-state level of the cIV subunits most profoundly decreased in 6B1^KO^ (MT-CO1, MT-CO2, COX6C) (Figures 4A, S3A). In more detail, LFQ-MS analysis of 6B1^KO^+AOX compared to 6B1^KO^ showed a remarkable increase of the MT-CO2 module subunits (MT-CO2, COX5B and COX7C), whereas the rest of cIV subunits showed a milder increase after AOX expression (Figure 4B). Interestingly, the proteomic analysis revealed a major increase in PET100, which was the most downregulated cIV AF in 6B1^KO^ compared to wt (Figure 1D). In addition, the AFs COX19, COX11 and MR-1S were also increased in 6B1^KO^+AOX (Figure 4B). In accordance, 2D BN/SDS-PAGE followed by WB showed increased incorporation of cIV subunits (MT-CO1, COX4, COX5A, MT-CO2, COX6C) into the assembly intermediate IV_sub_ in 6B1^KO^+AOX compared to 6B1^KO^, indicating that AOX-mediated alleviation of the reductive stress associated with cIV deficiency allows cIV assembly to proceed, specifically permitting the incorporation of MT-CO2 (Figures 4C, 4D, S3B). The abundance of both MT-CO1 and all the other tested cIV subunits in SC I III_2_IV_sub_ was also substantially augmented after AOX expression compared to 6B1^KO^, while IV_2_ and III_2_IV SCs remained absent (Figures 4C, 4D, S3B). Next, we examined whether the cIV forms assembled in 6B1^KO^+AOX were respiratory competent. Even though the AOX-expressing cells displayed robust rates of routine respiration (Figure S3C), titration of the AOX inhibitor (SHAM) completely abolished oxygen consumption, indicating that in intact cells oxygen consumption was exclusively happening through AOX. Further, supplementation of respiratory substrates into digitonin-permeabilized 6B1^KO^+AOX could not sustain significant oxygen consumption, again indicating an almost complete lack of respiration through cIV in these cells (Figure 4E). However, a low, yet still detectable respiration through cIV (COX) after supplementation with the artificial substrates ascorbate + TMPD (corrected for the KCN-insensitive portion) was observed in 6B1^KO^+AOX (Figure S3C), contrasting with 6B1^KO^ (Figure 4F). This means, that IV_sub_ present in 6B1^KO^ is catalytically competent, yet with only very limited activity, accounting only for approximately 5 % of the rate in wt cells.

**Figure 4:**
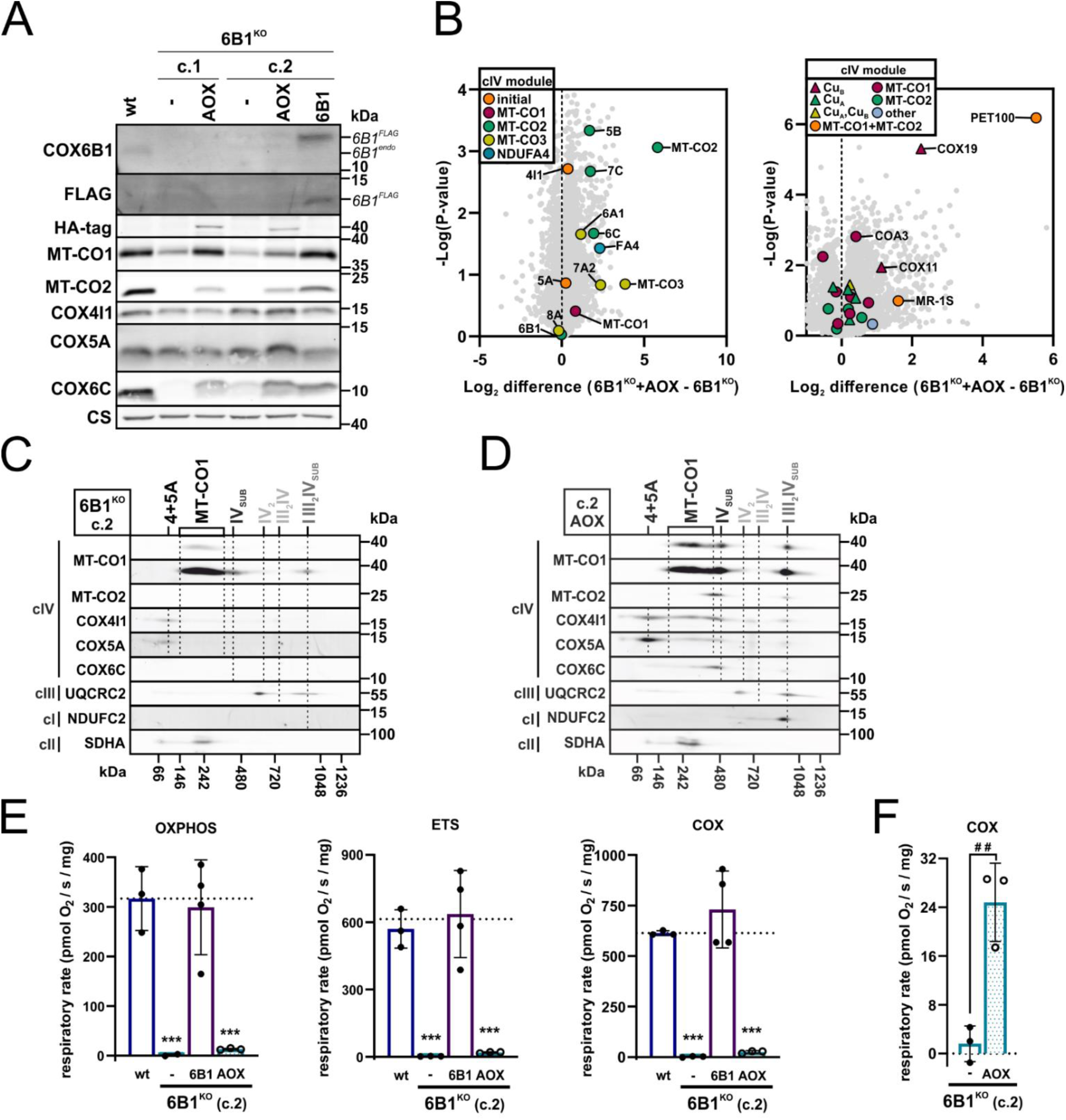
Expression of alternative oxidase restores cIV assembly intermediate formation with negligible cIV function in COX6B1 deficient cells. (A) Representative SDS-PAGE/WB analysis of the protein steady-state level of cIV subunits (MT-CO1, MT-CO2, COX4I1, COX5A, COX6B1, COX6C), C-terminal FLAG-tagged COX6B1 (FLAG antibody), HA-tagged AOX (HA-tag antibody), and CS as the loading control in whole cell-lysates. FLAG-tagged COX6B1 and endogenous COX6B1 proteins are marked with 6B1^FLAG^ and 6B1^endo^ (in italics) respectively. See also Figure S3A. (B) Differential content of cIV subunits and AFs between 6B1^KO^ and 6B1^KO^+AOX cells. Volcano plots represent LFQ-MS analysis (6B1^KO^ c.2: n = 3, 6B1^KO^+AOX: n = 3) of all analyzed proteins (grey), and subunits of cIV modules (initial: orange, MT-CO1: magenta, MT-CO2: green, MT-CO3: yellow, NDUFA4: blue) on the left, cIV assembly factors (MT-CO1 metalation: triangle in magenta, MT-CO1 maturation: circle in magenta, MT-CO2 metalation: triangle in green, MT-CO2 maturation: circle in green, MT-CO1+MT-CO2 metalation: triangle in yellow, MT-CO1+MT-CO2 association: circle in orange, other: circle in blue) on the right. COX6B1 protein missing in 6B1^KO^ was visualized thanks to the imputation step (missing values replaced from normal distribution) performed during the Perseus analysis of the LFQ-MS data. (C) 2D (BN/SDS)-PAGE/WB detection of cIV (MT-CO1, MT-CO2, COX4I1, COX5A, and COX6C antibodies), cIII (UQCRC2 antibody), and cI (NDUFC2 antibody) in 6B1^KO^ c.2 mitochondrial fraction. Antibody against cII (SDHA) was used as a loading control. See also Figure S3B. (D) 2D (BN/SDS)-PAGE/WB detection of cIV (MT-CO1, MT-CO2, COX4I1, COX5A, and COX6C antibodies), cIII (UQCRC2 antibody), and cI (NDUFC2 antibody) in 6B1^KO^+AOX mitochondrial fraction. Antibody against cII (SDHA) was used as a loading control. See also Figure S3B. (E) Respiratory rates in OXPHOS, ETS, and COX states plotted as the mean ± S.D. value of wt (n = 3), 6B1^KO^ (n = 3), 6B1^KO^+6B1 (n = 4) and 6B1^KO^+AOX (n = 3). One-way ANOVA (*** p < 0.001) was performed. Plotted data of wt, 6B1^KO^ and 6B1^KO^+6B1 originate from Figure 1G. (F) Respiratory capacity in COX state plotted as the mean ± S.D. value of 6B1^KO^ (n = 3) and 6B1^KO^+AOX (n = 3). Paired t test (# # p < 0.01) was performed. The graph represents y-axis zoomed-in representation of 6B1^KO^ and 6B1^KO^+AOX rates displayed in Figure 4E. See also Figure S3C.

To characterize the precise composition of the accumulated cIV assembly intermediates and the partial respirasome (SC I III_2_IV_sub_) in parental 6B1^KO^ cells and those expressing AOX, we performed proteomics-based complexome profiling analysis ^43^ (Figures 5A, 5B, S4A). Wt cells showed uniform distribution of the thirteen detected cIV subunits into all the mature cIV-containing species, whereas in 6B1^KO^ eight early-assembling subunits mostly accumulated in assembly intermediates (Figure 5A). Interestingly, a number of cIV subunits (COX4I1, MT-CO2, COX7A2) were detected within the SC I III_2_IV_sub_ in the 6B1^KO^ samples by complexome profiling. This indicates the incorporation of partially assembled cIV modules directly on the supercomplex platform running in parallel with the assembly of the individual complex, which is consistent with the ‘cooperative assembly’ model of respiratory chain biogenesis ^22^. In 6B1^KO^ cells expressing AOX, additional cIV subunits (including MT-CO3) were present in higher quantities in the SC I III_2_IV_sub_, only lacking a few late assembling cIV subunits (COX7C, COX8A, COX6A and NDUFA4), apart from the genetically removed COX6B1 (Figure 5A, 5B). In addition to the early assembly intermediates, the cIV subassembly (IV_sub_) containing almost all cIV subunits was also present in 6B1^KO^+AOX. Notably, the subunits detected in IV_sub_ belong to all three assembly modules and the only ones lacking were COX8A, COX6A and NDUFA4 (Figure 5A). These observations indicate that both the IV_sub_ and the SC I III_2_IV_sub_ represent the genuine and the most complex cIV species that can be assembled in the absence of COX6B1 in redox conditions that circumvent the assembly block associated with the absence of this particular subunit.

**Figure 5:**
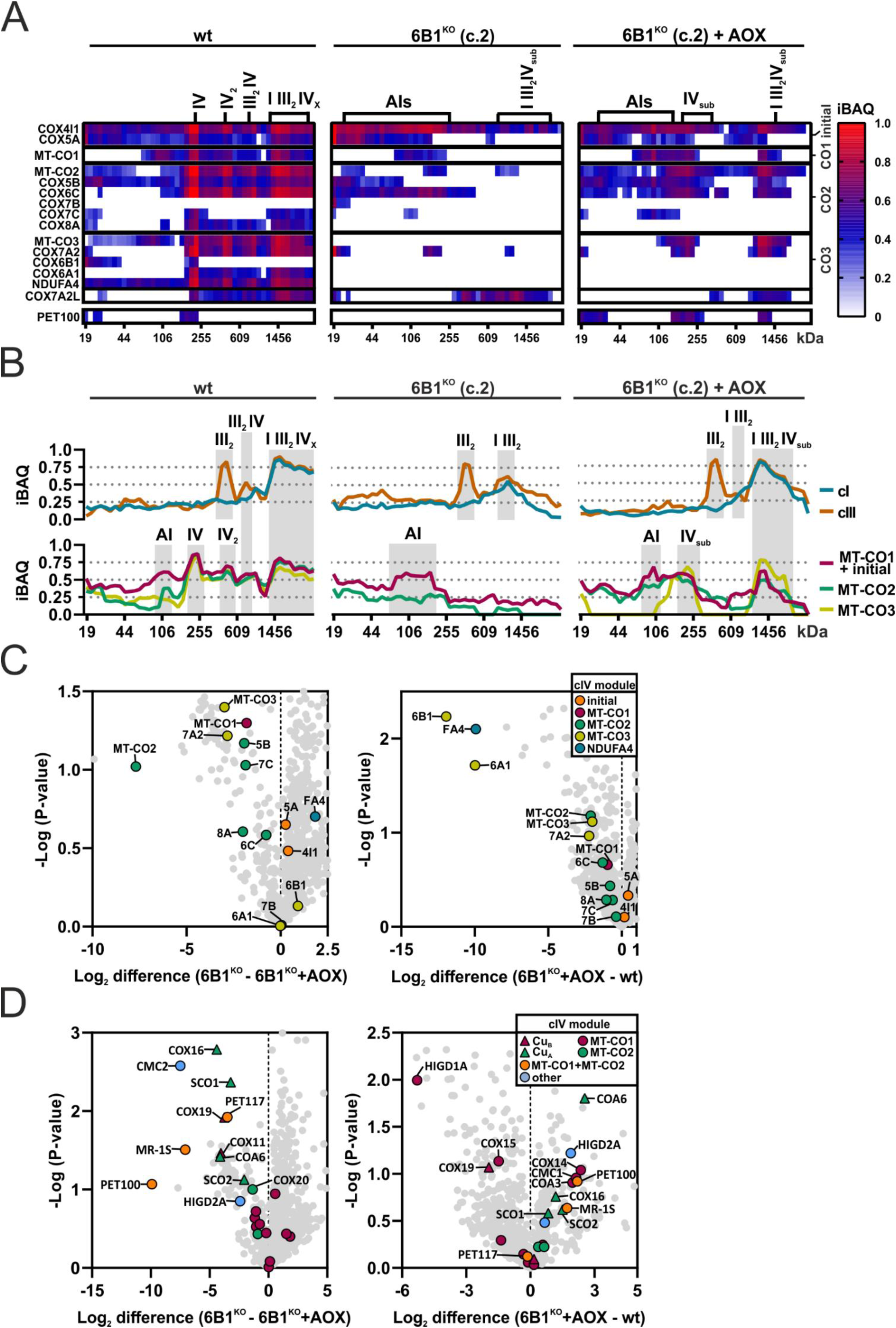
Expression of alternative oxidase restores early cIV assembly block in COX6B1 deficient cells. (A) Complexome profiling analysis of cIV in wt, 6B1^KO^, and 6B1^KO^+AOX respectively. Heat-map representation of relative iBAQ values of detected subunits of cIV (individual modules of are marked on the right) and AF PET100. (B) Complexome profiling analysis of cI, cIII and cIV in wt, 6B1^KO^, and 6B1^KO^+AOX respectively. XY graphs represent an average of relative iBAQ values of individual subunits of cI (blue) and cIII (orange) detected (top). In the case of cIV (bottom), average of subunits corresponding to the individual modules (initial + MT-CO1 in magenta, MT-CO2 in green, MT-CO3 in yellow) was made. In the case of MT-CO3 module in 6B1^KO^, no data were applied, since MT-CO3 subunit was not detected. See also Figure S4A. (C) Differential amount of cIV subunits associated in cIV forms between 6B1^KO^ and 6B1^KO^+AOX (left), 6B1^KO^+AOX and wt (right). Volcano plots represent LFQ-MS analysis (n = 2) of all detected proteins (grey), and subunits of cIV modules (initial: orange, MT-CO1: magenta, MT-CO2: green, MT-CO3: yellow, NDUFA4: blue). COX6B1 protein missing in 6B1^KO^ was visualized thanks to the imputation step (missing values replaced from normal distribution) performed during the Perseus analysis of the LFQ-MS data. (D) Differential amount of cIV AFs associated with cIV forms between 6B1^KO^ and 6B1^KO^+AOX (left), 6B1^KO^+AOX and wt (right). Volcano plots represent LFQ-MS analysis (n = 2) of all detected proteins (grey), and cIV assembly factors (MT-CO1 metalation: triangle in magenta, MT-CO1 maturation: circle in magenta, MT-CO2 metalation: triangle in green, MT-CO2 maturation: circle in green, MT-CO1+MT-CO2 metalation: triangle in yellow, MT-CO1+MT-CO2 association: circle in orange, other: circle in blue).

**Figure 6:**
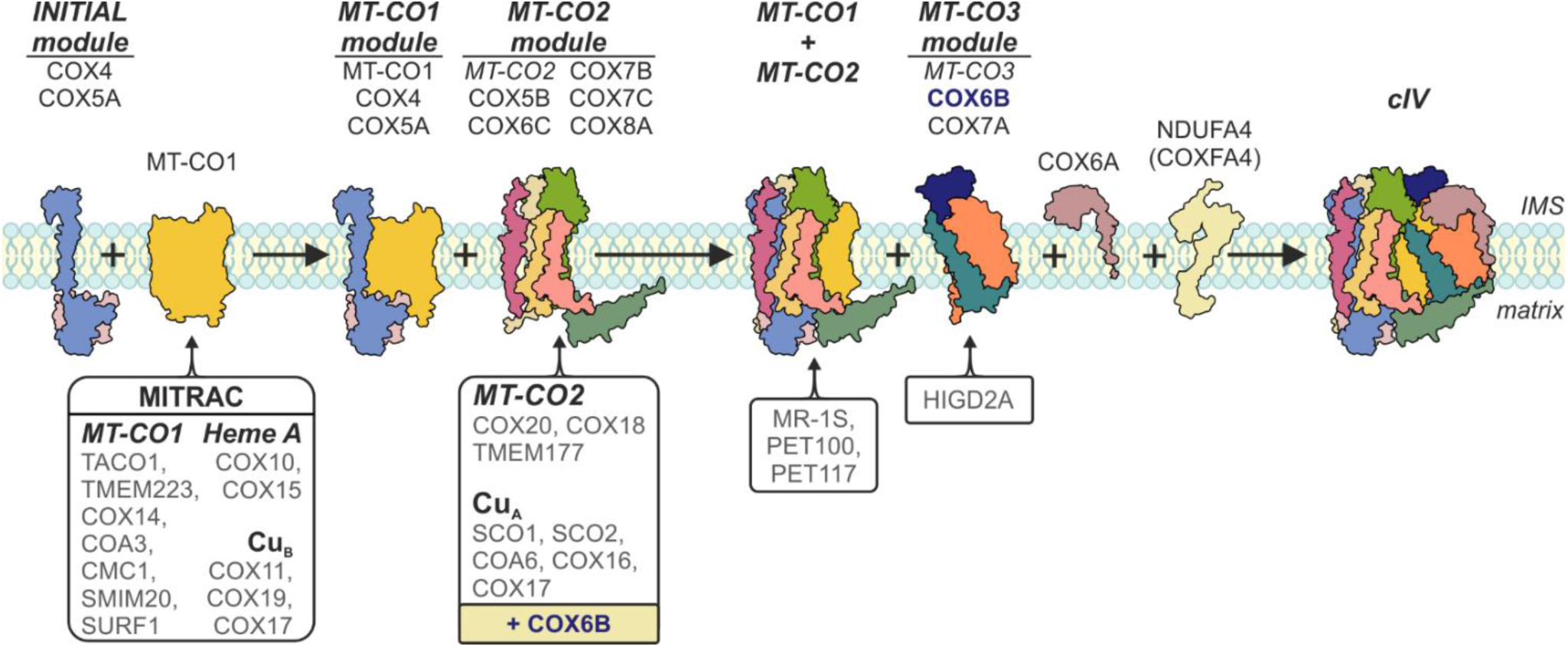
Updated modular model of human cIV assembly. The present scheme reflects the novel findings presented by this work. It implies COX6B subunit involvement in MT-CO2 subunit maturation. Also, COX6A subunit incorporates at the final assembly stages independent of the MT-CO3 module.

The complexome profiling results correlated with MS analysis of immunocaptured cIV, which showed a higher level of MT-CO2 and MT-CO3 module subunits bound to the native cIV forms in 6B1^KO^+AOX relative to 6B1^KO^, with the exception of COX6A1 and NDUFA4 subunits (Figure 5C). Moreover, another sign of restored MT-CO2 maturation and assembly after AOX expression in 6B1^KO^ was represented by augmented association of redox-sensitive AFs governing copper delivery to MT-CO2, i.e., SCO1, SCO2, COA6 and COX16, as well of the chaperones involved in MT-CO2 module incorporation (PET100, PET117, MR-1S) (Figure 5D). Interestingly, COA6, PET100 and other AFs were more abundant in 6B1^KO^+AOX than in wt, indicating persisting non-optimal assembly conditions in the absence of COX6B1, despite redox balance normalization by AOX (Figure 5D). This was specifically observed in the case of PET100 by complexome profiling, which was absent in 6B1^KO^, accumulated in the cIV monomer in wt, but was readily accumulated within IV_sub_ and I III_2_IV_sub_ in 6B1^KO^+AOX (Figure 5A). In addition, very low amounts of COX6A and NDUFA4 were immunocaptured with cIV in 6B1^KO^ compared to the wt (Figure 5C), consistent with the complexome profiling findings (Figure 5A), pointing out that the absence of COX6B1 allows the association of an incomplete MT-CO3 module lacking COX6A (Figure S4B).

## 3 Discussion

Complex IV deficiency is one of the most prominent causes of mitochondrial disease ^44,45^. The main genetic causes are pathogenic variants in nuclear genes encoding cIV assembly factors ^44,46^ and, less frequently, structural subunits ^8^. Despite the fact that proper cIV assembly is essential for its function and its implications for disease, many details of cIV biogenesis still remain unclear. This may be exemplified on COX6B1 subunit, which has always been recognized as a late-stage assembling subunit ^32^, being a part of the MT-CO3 module ^14^. Consistent with this idea, skin fibroblasts and muscle biopsies from patients carrying the COX6B1 R20C variant, showed an accumulation of a late cIV assembly intermediate ^29^. However, we have recently reported an unexpected total loss of cIV in HEK293 cell-line based COX6B1 knock-out (6B1^KO^), similarly to what was observed in a COX4 subunit KO model, lacking the initial module of cIV assembly ^20^. This observation argues against the peripheral role of COX6B in the maturation of human cIV.

In order to understand the real role of the COX6B subunit for cIV assembly, we have continued to investigate in detail the molecular mechanisms underlying the severe cIV deficiency caused by the complete absence of COX6B1. Thus, 6B1^KO^ cells showed decreased steady-state levels of most of the cIV subunits, including the mtDNA-encoded subunit MT-CO2. This was unexpected since, according to the current cIV assembly model, MT-CO2 precedes the addition of the MT-CO3 module, to which COX6B1 belongs ^14^. Unlike in the KO of COX4 subunit ^20,34^, 6B1^KO^ still preserved the initial module subunits COX4 and COX5A, reflected by the stabilization of assembly intermediates formed by subunits MT-CO1, COX4 and COX5A. In the 6B1^KO^ model, intermediates containing a residual portion of MT-CO2 subunit were present only in trace quantities detectable e.g. by complexome profiling, and a group of cIV subunits, i.e., COX4, MT-CO2, COX7A2 were found associated with a fully assembled SC I III_2_, forming an incomplete respirasome (SC I III_2_IV_sub_). However, the assembly intermediates migrating at a slightly lower molecular weight band than cIV monomer (IV_sub_), which would resemble the previous observation of incomplete cIV in COX6B1 patient samples (Massa et al 2008), were not detected in 6B1^KO^ cells, but were present when expressing the R20 COX6B1 variants. Therefore, we describe that COX6B1 subunit is indispensable for the addition or stabilization of the MT-CO2 module in an early phase of cIV assembly. In contrast, KO models of COX7A, another subunit belonging to the MT-CO3 module, did not affect the incorporation of MT-CO2 into cIV ^47^. Another evidence was a decrease in the steady-state and cIV-bound levels of the assembly factors governing copper delivery to MT-CO2 (SCO1, SCO2 and COA6), as well as MT-CO2 module stabilization and/or binding to the MT-CO1 module (PET100) in 6B1^KO^. Indeed, the phenotype of 6B1^KO^ mirrors COA6 lacking HEK293T cells ^37^, since both lead towards total cIV loss. Interestingly enough, COA6 is a paralog of COX6B1 ^48^ and their levels seem to be interdependent ^37^. Likewise, pathogenic variants in PET100 were shown to diminish MT-CO2 levels in patient fibroblasts ^49,50^, accompanied by a decrease in COX6B1 protein ^14^, leading to a total loss of fully assembled cIV ^14,50^.

The process of MT-CO2 metalation, i.e. Cu*_A_* insertion, requires intermembrane space located and redox-modulated assembly factors SCO1 and SCO2 and COA6, the latter containing twin Cx9C motifs involved in the redox-mediated import, folding and stabilization of the protein ^40,51^. COX6B1 also contains Cx9C (C30/C40) and Cx10C (C54/C65) motifs ^52^, which by analogy to COA6 and to the yeast COX6B1 orthologue Cox12, points out to a role for COX6B1 in MT-CO2 maturation ^53,54^. Xenoexpression of an alternative oxidase in 6B1^KO^ partially restored oxygen consumption, i.e., AOX-dependent electron flux through the respiratory chain, and this was associated with increased levels of most cIV subunits, with MT-CO2 level being the most significantly upregulated ^20^. Strikingly, cIV subunits detected across all the three assembly modules were stabilized in a cIV intermediate (IV_sub_), most likely as a result of ameliorated mitochondrial redox status, restoring MT-CO2 maturation. Accordingly, the factors required for coordination of the copper centers Cu*_A_* and Cu*_B_* ^51^ were enriched within the native forms of cIV after AOX expression compared with the parental 6B1^KO^. In accordance, blocked MT-CO2 metalation in 6B1^KO^ could also be inferred by the accumulation of COX14, CMC1 and COA3 within the immunocaptured cIV, representing a complex formed together with MT-CO1 that precedes the addition of MT-CO2 ^51^. In addition, the most significantly increased cIV AF in 6B1^KO^ after AOX expression was PET100, both at the steady-state level and cIV bound, indicating enhanced association of MT-CO1 and MT-CO2 modules. Therefore, here we present the first evidences that human COX6B1 subunit is instrumental for MT-CO2 maturation. So far, COX6B1 was related with this process only indirectly via a high-throughput BioID analysis of human cells using the SCO1 protein as a prey, where COX6B1 was fished out among SCO2, COA6, COX11, COX15, COX16, COX17, COX19 and COX5B ^55^.

Complexome profiling analysis of 6B1^KO^ cells expressing AOX, showed that incomplete cIV assembly modules, i.e., the MT-CO2 module missing COX7B, COX7C, and COX8A and the MT-CO3 module missing COX6A1 and COX6B1, assemble into SC I III_2_IV_sub_, while cIV_2_ and SC III_2_IV did not form. The absence of the cIV dimer may be explained by the proposed role of COX6B1 in cIV dimerization ^56,57^. Our findings also suggest that COX6A1 subunit is not able to join cIV in the absence of COX6B1, thus, representing the penultimate subunit of cIV assembly process preceding the last assembling subunit NDUFA4. Despite the absence of COX6B1 and a couple of other nuclear-encoded subunits, IV_sub_ or the supercomplex I III_2_IV_sub_ containing both MT-CO1 and MT-CO2 could theoretically provide complete electron-transfer pathway enabling enzyme activity. Our respirometric measurements demonstrated that the present IV_sub_ may indeed have enzymatic activity, yet it remains negligible. This finding supports the claim that COX6B1 though its extensive interactions with MT-CO2 is essential for the structure of the cytochrome *c* binding site ^58^. Alternatively, the impaired activity may result from incomplete maturation of the Cu*_A_* prosthetic factor in MT-CO2 in the absence of COX6B1 due to copper relay dysfunction ^54^. If this was the case, it could be speculated that modulation of the redox status in 6B1^KO^ by AOX only allows to pass a putative MT-CO2 maturation checkpoint in cIV assembly. This also opens the possibility that in this case, COX assembly proceeds through a non-canonical pathway that does not occur under normal circumstances. Future studies in cellular models with absence of other late-assembling subunits (COX6A, NDUFA4) should clarify these issues.

In a previous study, we reported the presence of the cI assembly intermediate without the catalytic N-module in contact with cIII dimer in COX4 lacking cells ^34^, indicating, also in this model, an association of partially formed individual respiratory chain complexes preceding their complete maturation. In 6B1^KO^ fully assembled SC I III_2_ was found associated with cIV subunits COX4-COX5A (pre-I III_2_IV_sub_), consistent with the previous observations of non-canonical *de novo* assembly of cIV directly into the nascent respirasome ^12,19^. In addition, PET100 was detected migrating not only with IV_sub_, but also along I III_2_IV_sub_ in 6B1^KO^+AOX, indicating active cIV assembly both in monomeric form and inside the respirasome. According to the present results, the addition of cIV subunits seems to be supported by the presence of an initial assembly intermediate formed by subunits MT-CO1, COX4, and COX5A, since the MT-CO1 signal was detected in the corresponding band only in 6B1^KO^ after AOX expression, but not in 4dKO ^20^. However, the possibility of some species of SC I III_2_IV_sub_ existing in 4dKO cells expressing AOX should be further explored, since MT-CO1 together with cIV subunits COX5B and COX7A2 was previously detected within SC I III_2_IV_sub_ in MT-CO2 lacking cells ^19^.

To date, the outcome of the COX6B1 pathogenic variants R20C and R20H was directly compared only in yeast models modelling the human variants in the *S. cerevisiae* orthologue *COX12* ^59^. Nevertheless, the applicability of these findings to molecular pathological mechanisms in humans may be limited due to the differences in the cIV structure and biogenesis between yeast and humans ^60^. Also, despite the impairment of the respiratory growth of the *COX12* null yeast cells, Cox2p levels were still preserved at around 50% of the wild type, and the introduction of R17H and R17C variants in Cox12p (analogous to R20H and R20C, respectively) did not complement the respiratory defect ^59^. Here, we have used novel HEK293 human cellular models with a stable expression of either R20C or R20H pathogenic variant of COX6B1 in the KO background. 6B1^KO^ phenotype improvement was limited in both R20C and R20H models, with partially increased respiratory rates. The observed differential extent of the cIV impairment due to R20C or R20H was originally explained by a destabilization of COX6B1 interaction within cIV, which is more pronounced in the case of R20C variant ^30^. However, this reasoning was based on the currently outdated structure of the bovine cIV ^61^, which did not contain the fourteenth cIV subunit NDUFA4 that is located in close proximity to R20 ^62^. Our findings imply that the additional Cys residue forming an artificial Cx9C motif of R20C may lead to aberrant folding and decreased content of the subunit, and possibly limits the MT-CO2 maturation and thus presents as a more severe cIV aberration compared to R20H variant. R20H variant rather destabilizes the association of COX6B1 with the complex but does not affect early cIV assembly to such an extent.

In this work, we show the dual role of COX6B1 subunit in the assembly of cIV. Nevertheless, the exact molecular mechanism by which COX6B1 interferes with other factors like PET100 and COA6, still requires further investigation. The exact role of the Cys residues of COX6B1 remains to be addressed, with an emphasis on the R20C pathogenic variant that represents a convenient model to clarify COX6B1 engagement in the MT-CO2 metalation. Also, since the COX6B2 isoform did not rescue the COX6B1 lacking HEK293 cell line, possibly for a tissue-/cell line-specific mechanism missing in these cells, its properties within the cIV biogenesis, structure and function will require a different study model than HEK293 cell line, i.e. spermatid cell line, sperm specimens or animal models. Finally, the role of individual nDNA-encoded subunits in the assembly process of monomeric cIV, as well as respiratory chain supercomplexes, should be revised in diverse knock-out models of cIV subunits and assembly factors for complete elucidation of the process, as demonstrated in this study by misconceptions of the role of COX6A and COX6B subunits.

## 4 Acknowledgements

The authors would like to acknowledge the Proteomics Service Laboratory at the Institute of Physiology (supported by RVO, ID 67985823) and the Institute of Molecular Genetics (supported by RVO, ID 68378050) of the Czech Academy of Sciences.

This research was supported by Czech Science Foundation projects (22-21082S, and 22-21552S), Czech Health Research Council (NU22-01-00499), and by the National Institute for Research of Metabolic and Cardiovascular Diseases (Program EXCELES, ID Project No. LX22NPO5104) funded by the European Union – Next Generation EU. K.C. is the recipient of an EMBO Postdoctoral Fellowship (ALTF 710-2022).

## Author contributions

Conceptualization, K.C., P.P.; Methodology, K.C., M.V., P.P.; Investigation, K.C., M.V., M.K., A.P., T.M., P.P.; Writing – Original Draft, K.C., P.P.; Writing – Review & Editing, K.C., A.P., L.A., J.H., E.F.V., T.M., P.P.; Visualization, K.C.; Funding Acquisition, P.P.; Supervision, T.M., P.P.

## 6 STAR methods

### 6.1 Resource availability Lead contact

Further information and requests for resources and reagents should be directed to and will be fulfilled by the lead contact, Petr Pecina.

#### Materials availability

- This study did not generate new unique reagents.

#### Data and code availability

- The mass spectrometry proteomics data from the label-free quantification (LFQ) analysis have been deposited to the ProteomeXchange Consortium via the PRIDE ^71^ partner repository with the dataset identifier PXD061621.
- The mass spectrometry proteomics data from the complexome profiling analysis have been deposited to the ProteomeXchange Consortium via the PRIDE ^71^ partner repository with the dataset identifier PXD061618.
- Original western blot images reported in this paper will be shared by the lead contact upon request.
- Any additional information required to reanalyze the data reported in this paper is available from the lead contact upon request.
- This paper does not report original code.

### 6.2 Experimental model and study participant detail

HEK293 (ATCC® CRL-1573™) cells, a cell line with epithelial morphology isolated from the kidney of a human embryo, were maintained at 37 °C and 5% CO_2_ atmosphere in DMEM/F-12 medium (Biowest, L0092) supplemented with 10% (v/v) FBS (Thermo Fisher Scientific, 10270-106), 40 mM HEPES, antibiotics (100 U/mL penicillin + 100 μg/mL streptomycin, Thermo Fisher Scientific, 15140-122) and 50 µM uridine.

### 6.3 Method details

#### Generation of HEK293 cellular models

*COX6B1* HEK293 (ATCC® CRL-1573™) knock-out (KO) cells (6B1^KO^, clone 1 – c.1, 2 – c.2, and 3 – c.3) were generated previously ^20^. For alternative oxidase (AOX) expression in 6B1^KO^ (c.1 and c.2) cells (to produce 6B1^KO^+AOX), pcDNA3.1+ mammalian expression vector (Thermo Fisher Scientific, USA) containing the coding sequence of full-length AOX (from *Aspergillus nidulans),* followed by a C-terminal HA-tag ^72^, was transfected using Metafectene Pro (Biontex Laboratories GmbH). For COX6B1 and COX6B2 expression in 6B1^KO^ (c.2) cells, pcDNA3.1^+^ mammalian expression vector (Thermo Fisher Scientific, USA) containing the coding sequence of full-length human COX6B1 or COX6B2 protein followed by a C-terminal FLAG-tag (to produce 6B1^KO^+6B1, and 6B1^KO^+6B2^C-FLAG^ respectively), or an N-terminal FLAG-tag in the case of COX6B2 (to produce 6B1^KO^+6B2^N-FLAG^), was transfected using Metafectene Pro (Biontex Laboratories GmbH). The pcDNA3.1^+^ construct with C-terminal FLAG-tagged COX6B1 was used to introduce mutations leading to single amino acid (AA) substitutions of arginine residue at position 20 (COX6B1-R20) to mimic two pathogenic variants of COX6B1 found in patients ^29,30^, either to cysteine (R20C) or histidine (R20H). Mutations were introduced by the QuikChange Lightning kit (Agilent, Santa Clara, CA, USA), using primers (see key resources table) suggested by the manufacturer’s on-line primer design tool. Mutagenesis was confirmed by sequencing, and mutated constructs were used for knock-in into 6B1^KO^ (c.2) cells to produce 6B1^KO^+R20C and 6B1^KO^+R20H models. Stably transfected cells were selected with 2 mg/mL G418.

#### SDS-PAGE

Proteins separation under denaturing conditions was performed using tricine-sodium dodecyl sulfate polyacrylamide gel electrophoresis (SDS-PAGE). Protein samples were prepared from frozen cellular pellets as described ^73,74^. 20-30 µg of protein were separated on 12 % polyacrylamide gels using the Mini-PROTEAN III apparatus (Bio-Rad, USA). Experiments were performed at least three times to assess the statistical significance of the results.

#### Native and 2D electrophoresis

For the separation of native protein complexes, blue-native gel electrophoresis (BN-PAGE) ^75^ was performed. For 2D (BN/SDS)-PAGE and complexome profiling, mitochondrial pellets were isolated from freshly harvested cells by hypotonic shock followed by differential centrifugation ^76^ and solubilized using the mild detergent digitonin (6 g detergent / 1 g protein) to preserve supercomplex association. Final samples (30 µg protein) ^74,75^ were separated on a 4-13 % polyacrylamide gradient gel using the Mini-PROTEAN III apparatus (Bio-Rad). Afterwards, gel was cut in single lanes and either used for complexome profiling analysis or for a second dimension of 2D electrophoresis (SDS-PAGE) after 1 h incubation in dissociation solution (1% SDS, 1% 2-Mercaptoethanol).

For BN-PAGE separation (Figure S4B), mitochondrial-enriched fraction was obtained by digitonin treatment of the freshly harvested cellular pellets, which was further solubilized by n-dodecyl-β-D-maltoside (DDM) based on the protocol ^77^. Samples were separated on a precast commercial native gel (NativePAGE 3-12% Bis-Tris Gels, Thermo Fisher Scientific, Cat#BN1001BOX) as described ^77^.

#### Western blotting (WB) and immunodetection

Proteins separated by SDS-PAGE were transferred on polyvinylidene difluoride (PVDF) membranes (Immobilon FL 0,45 µm, Merck) by semi-dry electroblotting (0.8 mA/cm^2^, 1 hour) using a Transblot SD apparatus (Bio-Rad). BN-PAGE gels were transferred to PVDF membranes using Dunn’s carbonate buffer (10 mM NaHCO_3_, 3 mM Na_2_CO_3_), applying a constant voltage of 100 V at 4°C for 1 hour using a Mini Trans-Blot® Cell (Bio-Rad). Immunodetection was performed as described ^74^, using the primary and fluorescent secondary antibodies listed in the Key resources table. The resulting signals were analyzed and quantified by Image Lab software (Bio-Rad).

#### Label-free quantification mass spectrometry analysis

Proteomic analysis was performed by the Proteomics Service Laboratory at the Institute of Physiology and the Institute of Molecular Genetics of the Czech Academy of Sciences. Label-free quantification mass spectrometry analysis (LFQ-MS) of cell pellets was performed. Briefly, cellular pellets (100 µg of protein) were processed according to the protocol for in-solution trypsin digestion ^34^. About 1 µg of peptide digests were separated on a 50 cm C18 column using 2.5 h gradient elution and analyzed in a DDA mode on the Orbitrap Exploris 480 (Thermo Fisher Scientific) mass spectrometer. The resulting raw files were processed in MaxQuant (v. 1.5.3.28) ^78^ with the label-free quantification (LFQ) algorithm MaxLFQ ^79^. Imputation of missing values was performed in Perseus using default values (replaced from normal distribution, width 0.3, down shift 1.8). LFQ-MS data are available via ProteomeXchange with identifier PXD061621.

#### Immunoprecipitation and mass spectrometry analysis

Mitochondrial membranes solubilized with digitonin (6 g/g of protein) were immunoprecipitated using the Complex IV immunocapture kit (Abcam, ab109801). Proteomic analysis was performed by the Proteomics Service Laboratory at the Institute of Physiology and the Institute of Molecular Genetics of the Czech Academy of Sciences. Washed beads with immunocaptured complexes were digested “on beads” with trypsin using the sodium deoxycholate procedure as described in ^80^. Desalted peptide digests were separated on a 50 cm C18 column using a 1 h elution gradient and were analyzed on Orbitrap Exploris 480 MS. Resulting raw files were processed in MaxQuant (v. 1.6.6.0, maxquant.org) with the label-free relative quantification (LFQ) algorithm MaxLFQ ^79^. Analysis was performed in a duplicate. Imputation of missing values was performed in Perseus using default values (replaced from normal distribution, width 0.3, down shift 1.8). LFQ-MS data are available via ProteomeXchange with identifier PXD061621.

#### Complexome profiling

Isolated mitochondria from wt, 6B1^KO^ and 6B1^KO^+AOX cells were solubilized by digitonin (6 g detergent / 1 g protein) and samples for BN-PAGE separation were prepared as described earlier. 50 µg of finalized samples were subjected to 4 – 13 % polyacrylamide gradient gel using Mini-PROTEAN III apparatus (Bio-Rad). The gel was fixed (1 methanol: 1 dH_2_O; 5% acetic acid), washed in dH_2_O, and excised in 48 slices. Slices were cut to 1 mm^3^ cubes and transferred to a 96-well filter plate (Multiscreen Solvinert MSRLN0450, 0.45 µm pore size PTFE membrane, Merck Millipore). Proteomic analysis was performed by the Proteomics Service Laboratory at the Institute of Physiology and the Institute of Molecular Genetics of the Czech Academy of Sciences. In-gel digestion was performed according to the protocol ^81^ modified for a 96-well filter plate processed on a vacuum manifold. Briefly, reduction by DTT was followed by iodoacetamide alkylation, de-staining, and trypsin digestion. Tryptic peptides were extracted, dried, dissolved, desalted by 1-layer C18 StageTips, and transferred to a U-shaped 96-well plate suitable for the Ultimate 3000 nano UHPLC autosampler. Peptides were separated on a 15 cm C18 column using 30 min elution gradient and analyzed in a DDA mode on Orbitrap Exploris 480 MS. Raw files were processed in MaxQuant (v. 1.6.17.0). The iBAQ values of each protein detected in the individual slices were normalized to the protein with the highest intensity and were visualized in the heatmaps using Microsoft Excel and Prism 8 (GraphPad Software, La Jolla California USA). Complexome profiling data are available via ProteomeXchange with identifier PXD061618.

#### High-resolution respirometry

Mitochondrial respiration was measured at 37 °C as described in ^82^ using Oxygraph-2k (Oroboros, Innsbruck, Austria), essentially as described in ^83^. Freshly harvested cells (0.4 – 1.5 mg of protein) were suspended in 2 mL of MiR05 medium (0.5 mM EGTA, 3 mM MgCl_2_, 60 mM lactobionic acid, 20 mM taurine, 10 mM KH_2_PO_4_, 20 mM HEPES, 110 mM D-sucrose, and 1g/l of BSA, pH 7.1) ^84^, and digitonin (0.05 g/g protein) was used to permeabilize the plasma membrane. For measurements, the following substrates and inhibitors were used: 2 mM malate, 10 mM pyruvate, 10 mM glutamate, 10 mM succinate, 10 mM glycerol 3-phosphate, 1 mM ADP, 0.5 µM oligomycin, 0.5–2 µM FCCP, 0.5 µM rotenone, 10 mM malonate, 0.25 µM antimycin A, 2 mM ascorbate, 1 mM TMPD, 0.5 mM KCN, 0.5 mM salicylhydroxamic acid (SHAM, inhibitor of AOX). The oxygen consumption was expressed in pmol O_2_/s/mg protein. Data were acquired and analyzed using routine functions of Datlab 5 software (Oroboros). Representative trace of the respiratory protocol is shown in Figure S1E. Three respiratory states were chosen for data reporting: i) OXPHOS – actively phosphorylating cells supplied fully saturated with substrates (pyruvate, glutamate, malate, succinate and glycerol 3-phosphate) and ADP, ii) ETS – maximal respiratory capacity of the electron-transporting system (fully saturated with substrates, after FCCP uncoupling), iii) COX – oxygen consumption by cytochrome *c* oxidase fueled by artificial substrates ascorbate + TMPD, corrected for KCN-insensitive background oxygen consumption).

#### Data analysis, visualization, and statistics

Data were analyzed and visualized in GraphPad Prism 8 software (GraphPad Software). Downstream analysis of LFQ-MS data was performed in Perseus (v. 2.3.0.0., v. 2.0.11.0.) ^78^ and data were further visualized in GraphPad Prism 8. One-way ANOVA (Asterisks represent p value: * <0.05; ** <0.01; *** <0.001) was used. Significant changes between individual samples and wt are represented by a symbol over the corresponding bar, while comparison between other models is indicated by a star symbol on top of a connecting line. Data shown in the graphs represent the mean values ± S. D. of at least 3 independent experiments.

The key resource table is attached as individual supplemental file.

