## supplemental figures for "COX6B1 secures a redox-sensitive step in early cytochrome *c* oxidase assembly"

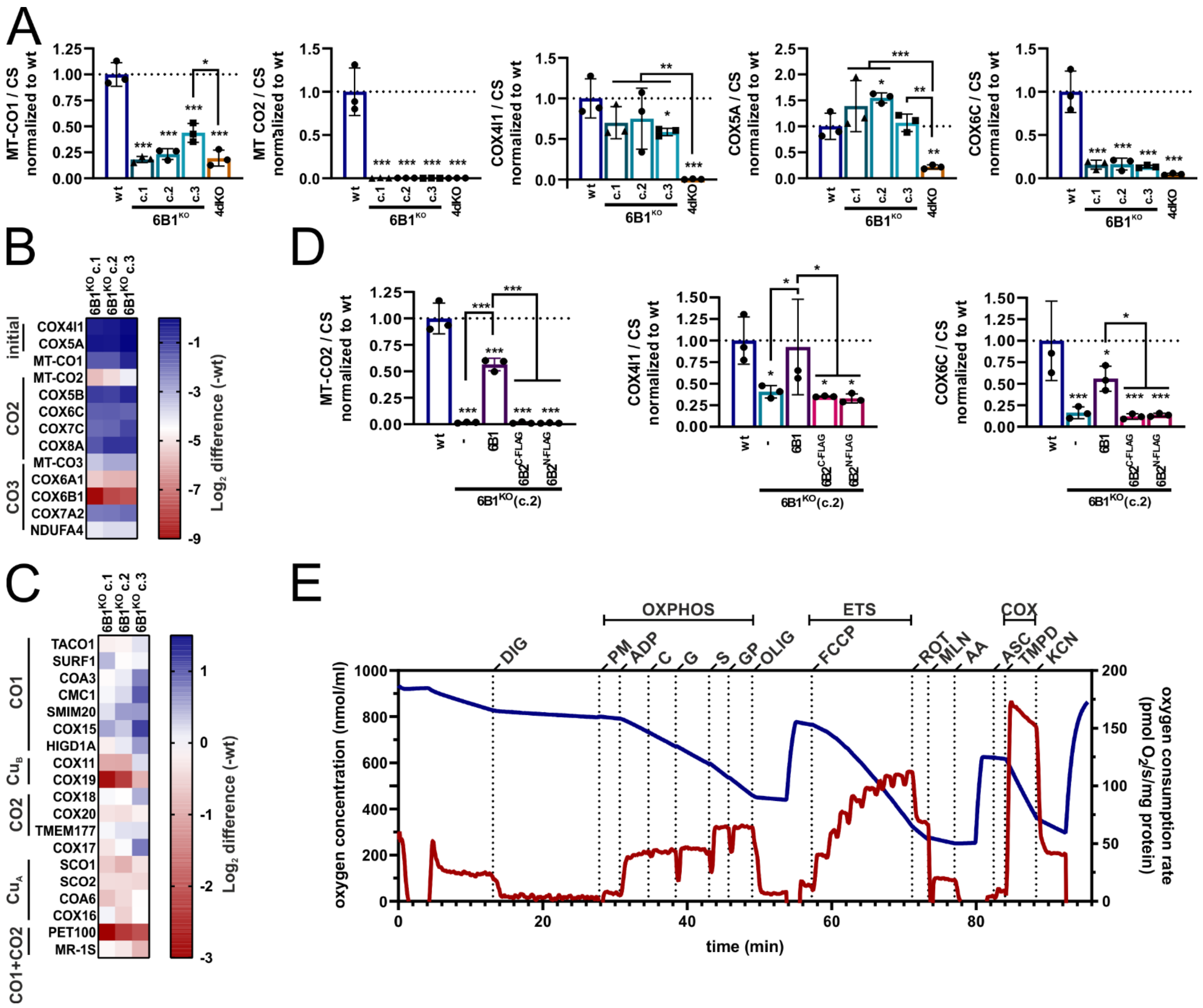

**Figure S1: COX6B1 knock-out blocks early human cIV assembly**

(A) Quantification of the MT-CO1, MT-CO2, COX4I1, COX5A, COX6C signals from SDS-PAGE/WB analysis normalized to CS. One-way ANOVA (* $p < 0.05$; ** $p < 0.01$; *** $p < 0.001$) was performed ($n = 3$, mean ± SD).

(B) Differential content of cIV subunits between wt and individual 6B1KO clones. Heatmap represents LFQ-MS analysis (wt: $n = 4$; 6B1KO c.1, c.2 and c.3: $n = 2$ per each) of analyzed subunits of cIV modules. COX6B1 protein missing in 6B1KO was visualized thanks to the imputation step performed during the Perseus analysis of the LFQ-MS data.

(C) Differential content of cIV assembly factors (AFs) between wt and individual 6B1KO clones. Heatmap represents LFQ-MS analysis (wt: $n = 4$; 6B1KO c.1, c.2 and c.3: $n = 2$ per each) of analyzed cIV assembly factors.

(D) Quantification of the MT-CO2, COX4I1 and COX6C signals from SDS-PAGE/WB analysis normalized to CS. One-way ANOVA (* $p < 0.05$; ** $p < 0.01$; *** $p < 0.001$) was performed ($n = 3$, mean ± SD).

(E) Representative trace of respirometric measurement of wt cells. Experimental trace recorded by Oxygraph-2k (Oroboros) shows the actual O2 concentration (blue, left Y axis) and rate of oxygen consumption (red, right Y axis). Additions of substrates and inhibitors are marked by vertical dashed lines and abbreviations above the trace (digitonin - DIG, pyruvate + malate - PM, ADP - ADP, cytochrome c - C, glutamate - G, succinate - S, glycerol-3 phosphate - GP, oligomycin - OLIG, FCCP - FCCP, rotenone - ROT, malonate - MLN, antimycin A - AA, ascorbate – A, TMPD - T, and KCN - KCN).

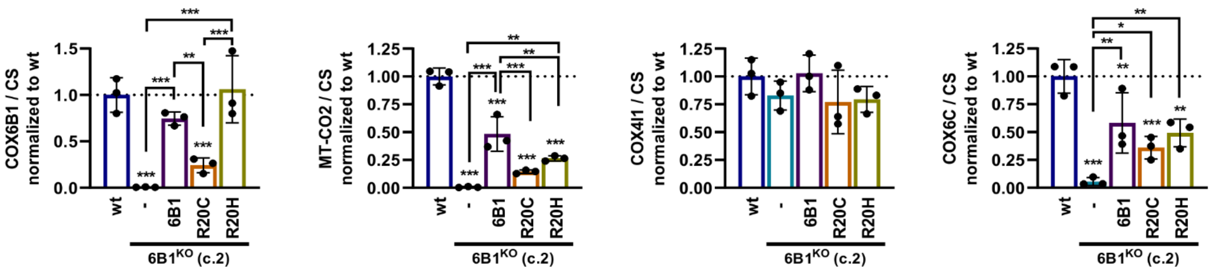


**Figure S2: R20C pathogenic variant of COX6B1 protein is less stable than R20H**
Quantification of the COX6B1, MT-CO2, COX4I1, COX6C signals from SDS-PAGE/WB analysis normalized to CS. One-way ANOVA (* $p < 0.05$; ** $p < 0.01$; *** $p < 0.001$) was performed ($n = 3$, mean ± SD).

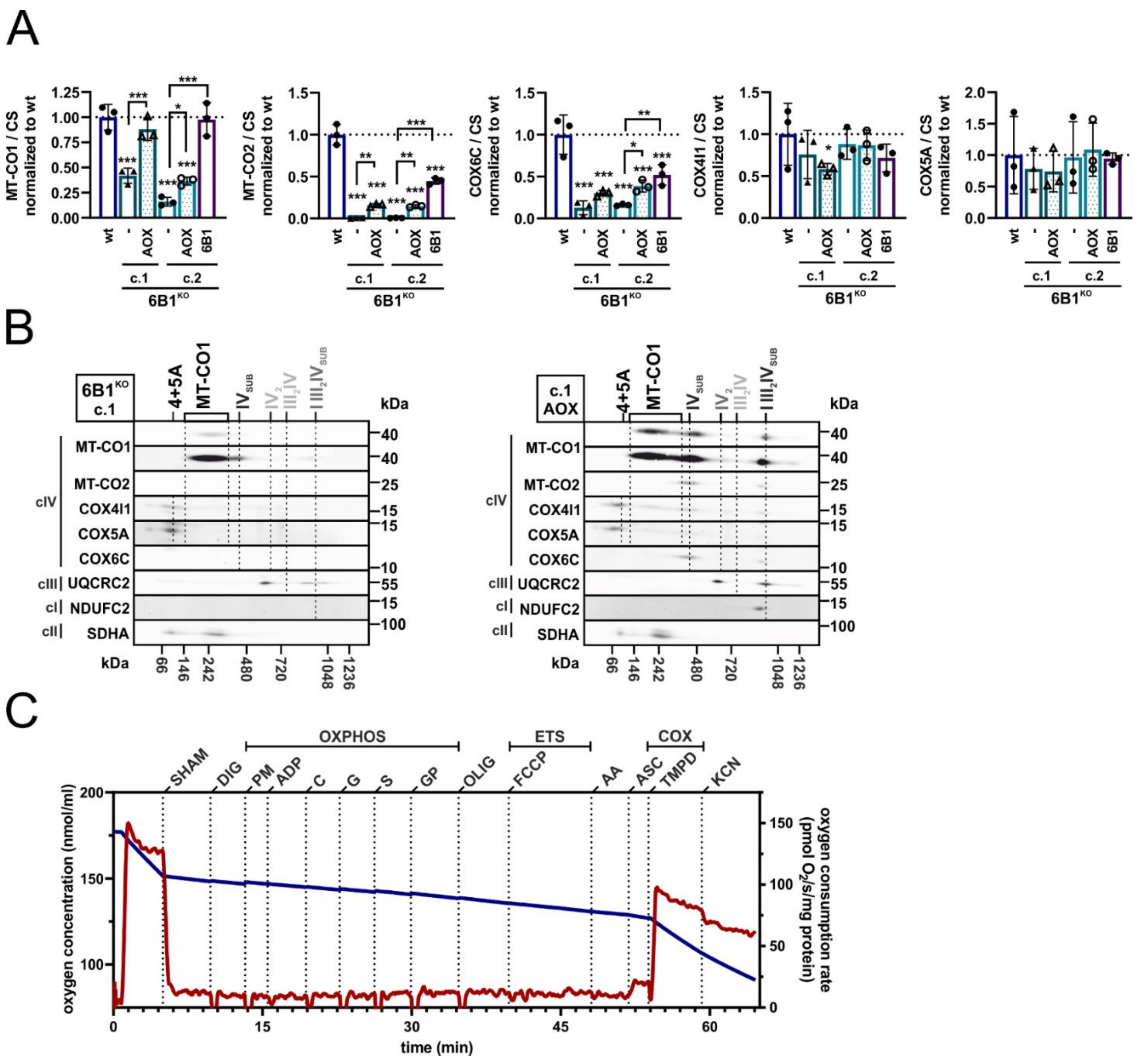


**Figure S3: Alternative oxidase expression ameliorates cIV composition and function in COX6B1 deficient cells**

(A) Quantification of the MT-CO1, MT-CO2, COX4I1, COX5A, COX6C signals from SDS-PAGE/WB analysis normalized to CS. One-way ANOVA (* $p < 0.05$; ** $p < 0.01$; *** $p < 0.001$) was performed ($n = 3$, mean ± SD).

(B) 2D (BN/SDS)-PAGE/WB detection of cIV (MT-CO1, MT-CO2, COX4I1, COX5A, and COX6C antibodies), cIII (UQCRC2 antibody), and cI (NDUFC2 antibody) in 6B1KO c.1 (left) and 6B1KO c.1 +AOX (right) mitochondrial fraction. Antibody against cII (SDHA) was used as a loading control.

(C) Representative trace of respirometric measurement of 6B1KO+AOX. Experimental trace recorded by Oxygraph-2k (Oroboros) shows the actual O2 concentration (blue, left Y axis) and rate of oxygen consumption (red, right Y axis). Additions of substrates and inhibitors are marked by vertical dashed lines and abbreviations above the trace (digitonin - DIG, pyruvate + malate - PM, ADP - ADP, cytochrome c - C, glutamate - G, succinate - S, glycerol-3 phosphate - GP, oligomycin - OLIG, FCCP - FCCP, antimycin A - AA, ascorbate – A, TMPD - T, and KCN - KCN).

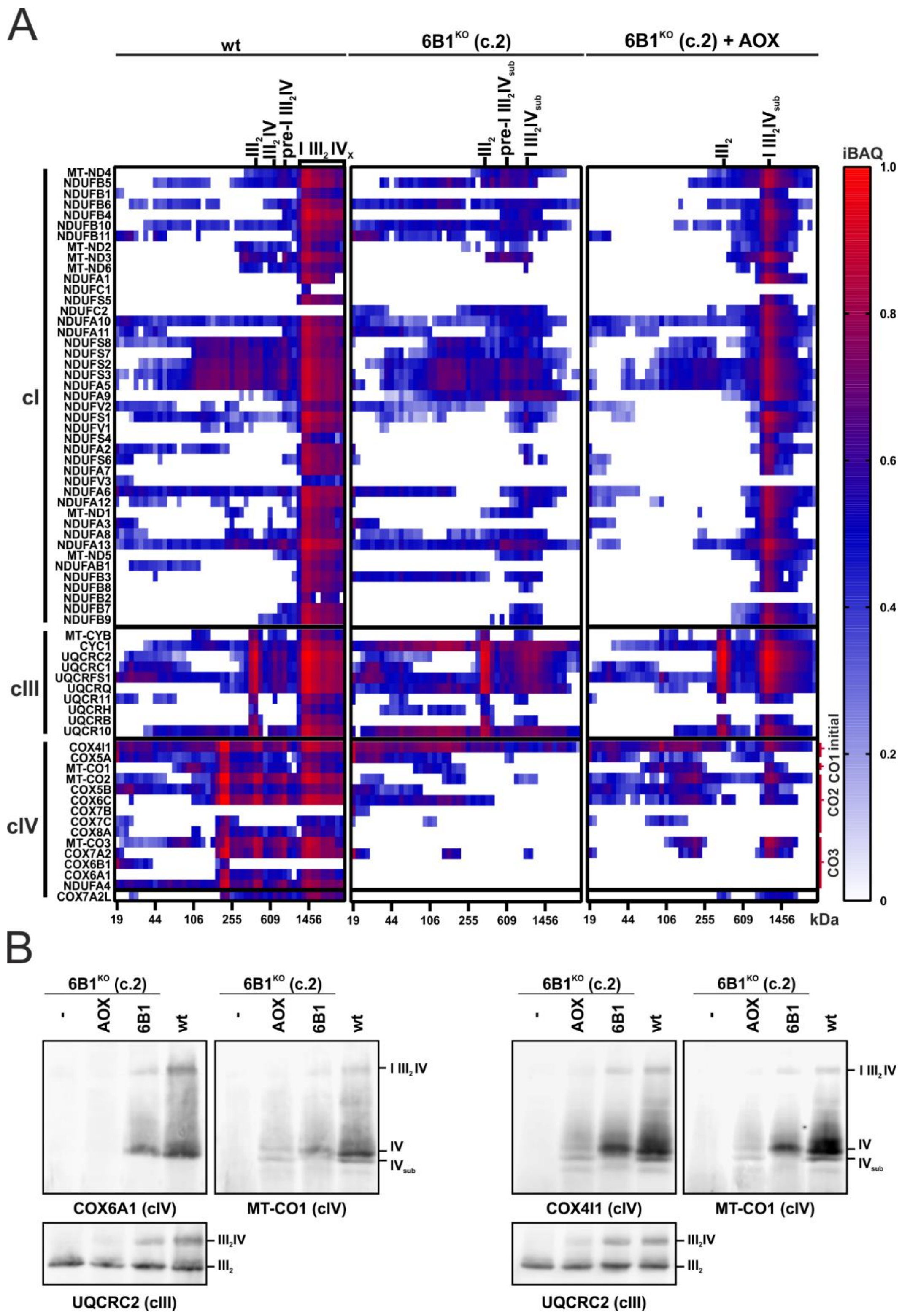


**Figure S4: Alternative oxidase expression restores cIV assembly in COX6B1 deficient cells**
(A) Complexome profiling analysis of cI, cIII and cIV subunits in wt, 6B1KO, and 6B1KO+AOX respectively. Heat-map representation of relative iBAQ values.
(B) Blue-native (BN)-PAGE/WB detection of cIV using COX6A1, MT-CO1 and COX4 antibodies. Antibody against cIII (UQCRC2) was used as a loading control.
